# CVB3 VP1 interacts with MAT1 to inhibit cell proliferation by interfering with Cdk-activating kinase complex activity in CVB3-induced acute pancreatitis

**DOI:** 10.1101/2020.09.21.305938

**Authors:** Hongxia Zhang, Lingbing Zeng, Qiong Liu, Guilin Jin, Jieyu Zhang, Zengbin Li, Yilian Xu, Xiaotian Huang

## Abstract

Coxsackievirus B3 (CVB3) belongs to the genus *Enterovirus* of the family *Picornaviridae* and can cause acute acinar pancreatitis in adults. However, the molecular mechanisms of pathogenesis underlying CVB3-induced acute pancreatitis have remained unclear. In this study, we discovered that CVB3 capsid protein VP1 inhibited pancreatic cell proliferation and exerted strong cytopathic effects on HPAC cells. Through yeast two-hybrid, co-immunoprecipitation, and confocal microscopy, we show that Menage a trois 1 (MAT1), a subunit of the Cdk-Activating Kinase (CAK) complex involved in cell proliferation and transcription, is a novel interaction protein with CVB3 VP1. Moreover, CVB3 VP1 inhibited MAT1 accumulation and localization, thus interfering with its interaction with CDK7. Furthermore, CVB3 VP1 could suppress CAK complex enzymic phosphorylation activity towards RNA Pol II and CDK4/6, direct substrates of CAK. VP1 also suppresses phosphorylation of retinoblastoma protein (pRb), an indirect CAK substrate, especially at phospho-pRb Ser^780^ and phospho-pRb Ser^807/811^ residues, which are associated with cell proliferation. Finally, we present evidence using deletion mutants that the C-terminal domain (VP1-D8, 768-859aa) is the minimal VP1 region required for its interaction with MAT1, and furthermore, VP1-D8 alone was sufficient to arrest cells in G1/S phase as observed during CVB3 infection. Taken together, we demonstrate that CVB3 VP1 can inhibit CAK complex assembly and activity through direct interaction with MAT1, to block MAT1-mediated CAK-CDK4/6-Rb signaling, and ultimately suppress cell proliferation in pancreatic cells. These findings substantially extend our basic understanding of CVB3-mediated pancreatitis, providing strong candidates for strategic therapeutic targeting.

## Introduction

Cholelithiasis and alcoholism are the most common risk factors of acute pancreatitis (AP). However, approximately 10% of AP occurs in patients with other miscellaneous evidence such as Coxsackie Virus group B infection. Coxsackie Virus group B3 (CVB3), in genus *Enterovirus* of family *Picornaviridae*, can cause a variety of human diseases, including myocarditis, pancreatitis, and can even lead to sudden infant death [1-7]. However, the mechanism by which CVB3 infection can cause pancreatitis remains unclear.

The crystal structure of CVB3 revealed the virus to be icosahedral, and without an envelope. CVB3 encodes four capsid proteins (VP1, VP2, VP3 and VP4), of which VP1 accumulates in the highest concentrations. Recent studies on the VP1 protein have primarily focused on vaccine development and its interactions with cell surface receptors. B-cell epitopes located on VP1 (VP1 1-15 aa, VP1 21-35 aa, and VP1 229-243 aa) and T-cell epitopes on VP1 (VP1 681–700 aa, VP1 721–740 aa, and VP1 771–790 aa) have been considered ideal vaccine candidates for protection against CVB3 infection[8, 9]. A VP1 protein nanoparticle vaccine and CPE30-chitosan-VP1 nanoparticles were both shown to prevent CVB3 induced myocarditis. In addition, interrogation of the CVB3-DAF (human decay-accelerating factor) complex structure, determined by cryo-electron microscopy, confirmed that DAF S104 is an essential site for close contact with VP1 residue T271 during CVB3 invasion[10-13]. Our previous work also identified Golgi Matrix Protein 130 (GM130) as a direct intracellular target of CVB3 VP1, and indicated that the interaction between VP1 and GM130 could disrupt the structure of the Golgi ribbon[14]. However, few studies have reported the effects of VP1 on cell proliferation and its direct protein targets in cells.

Cell proliferation is the process by which cells increase in number and is mediated by the core cell cycle machinery in response to various external and internal signals. Typically, the cell cycle is controlled by cyclin dependent kinases (CDKs). The activation of CDKs relies on the phosphorylation of essential threonine residues in the T - loop by CDK-activated kinase (CAK) [15]. The CAK complex is composed of MAT1, CDK7, and Cyclin H [16, 17] and is a subcomplex of TFIIH (Transcription Factor II H) which is comprised of ten subunits and is essential for transcriptional initiation. MAT1 is required for functional CAK assembly and induces CDK7 kinase activity[18]. However, TFIIH depends on CDK7 activity to play its role in transcriptional initiation[19, 20]. CDK7 has been revealed to perform dual functions, contributing a central role in the enzymatic activity of the CAK complex while also regulating transcription through phosphorylation of RNA polymerase II and transcription factors[8, 21, 22].

Other CDK family proteins, such as CDK2, CDK4, and CDK6, are involved in regulating cell cycle[23]. CDK2, CDK4/6, and RNA polymerase II are activated via direct phosphorylation by CAK. Activated CDK2 and CDK4/6 can then phosphorylate Rb (Retinoblastoma tumor suppressor protein), resulting in activation of E2F and release of transcription factors necessary for G1/S phase progression[24, 25]. Phosphorylation of pRb at several clusters of sites could inhibit E2F binding, especially at the C-terminal sites such as Ser780, Ser795 and Ser807/811[26]. The activated RNA polymerase II is then able to bind DNA and start transcription of S-phase genes[27-30]. The CDK-Rb-E2F pathway is undoubtedly responsible for G1 phase progression and transition to the S phase[25]. In this context, CDK2, CDK4/6, and RNA polymerase II can be understood as direct substrates of CAK, while Rb serves as an indirect substrate. Indeed, there were a few examples of viral modulation of cyclin activity to regulate cell proliferation. Avian Reovirus p17 protein is reported to suppress cell proliferation by directly binding to CDK-associated proteins, except for the CDK1-cyclin B1 and CDK7-cyclin H complexes[31]. Furthermore, pUL21a, a Human cytomegalovirus (HCMV) protein, was found to interact with cyclin A, causing the degradation of the proteasome through binding with its cyclin-binding domain[32]. HIV Tat protein was shown to target CDK9 as part of its contrubiton to HIV transcription [33, 34]. However, the molecular mechanisms by which CVB3 proteins directly regulate CDKs and influence cell proliferation are still unknown.

In this study, we show that structural protein CVB3 VP1 can reduce HPAC cell proliferation by arresting the cell cycle at the G1/S phase. More interestingly, by interacting with MAT1, VP1 impairs the structural formation and activity of the CAK complex via mediating CAK-CDK4/6 signaling pathway, which functions as a switch for cell cycle initiation in pancreatic cells, and also reduces CDK7 expression. Moreover, we also identified that VP1-D8 (198-284aa) was the minimum domain of VP1 as function to inhibit cell cycle in the G1 phase through the deletion mutants approach. These findings describe a previously unreported phenomenon of in which the CVB3 VP1 structural protein inhibits pancreatic cell proliferation by interfering with cell cycle through impairment of CAK complex assembly and function. This work thus modifies our understanding of the role of CVB3 in causing pancreatitis mechanisms and opens new avenues for targeted therapies.

## Results

### CVB3 capsid protein VP1 inhibits HPAC cell proliferation

First, to quantitatively determine the cytotoxicity of VP1 towards HPAC cells, we observed its cytopathic effects and performed adherent cell counts. We found that transfection of VP1 resulted in a reduction in the number of adherent HPAC cells (Fig 1A), we also observed that VP1 induced HPAC cell shrinking, detachment, and lysis (Supplemental Fig. 1). We then explored the effects of CVB3 VP1 on HPAC cell proliferation.

**Figure 1.**
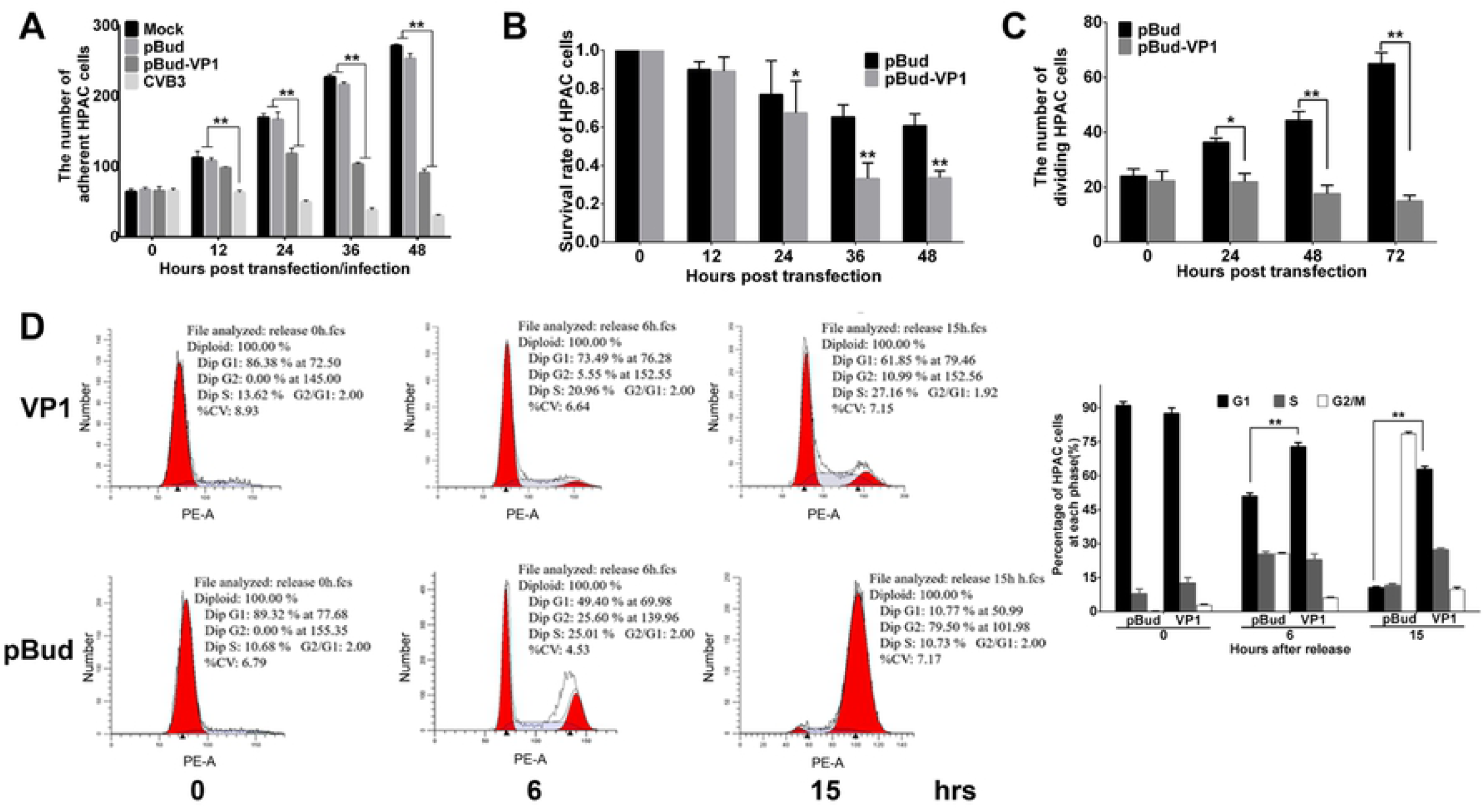
CVB3 VP1 affects cell proliferation and blocks cell cycle at the G1/S phase. **(A)** The numbers of adherent HPAC cells in different groups were counted using Image Pro Plus (IPP) software. **(B)** HPAC cells were transfected by pBud-VP1 and pBud plasmids, and the cell proliferation was evaluated using the CCK-8 assay. The activity of DNA replication was examined using EdU incorporation in HPAC cells. **(D)** G1/S phase arrest induced by VP1 transfection. HPAC cells were treated with 36-hour plasmid transfection (pBud and pBud-VP1) within the treatment of double-thymidine, and collected at 0, 6 and 15 hpi separately, then cells were analyzed by flow cytometry. Values are shown as the mean ± SEM of three independent experiments. (\**P*<0.05, \*\**P*<0.01).

Analysis of CCK-8 assays confirmed that VP1 reduced the cell survival rate compared with that of the vector control group with extension of transfection time in HPAC cells (Fig. 1B). EdU assays also showed that DNA replication was significantly inhibited in HPAC cells transfected with VP1, also commensurately with increasing time of transfection (Fig. 1C). Western blot analysis revealed that the VP1 expression decreased with the time course of VP1 transfection or CVB3 infection (Supplemental Fig. 2). Further, to characterize VP1 cytotoxicity towards HeLa cells, we quantified adherent, viable, and proliferating cells after exposure to VP1. We found that VP1 significantly inhibited HeLa cell proliferation, and showed the same cytotoxic effects as those observed in HPAC cultures (Supplemental Fig. 3). These results showed that the effects of VP1 alone was consistent with that of CVB3 virus infection in both HPAC and HeLa cells.

**Figure 2.**
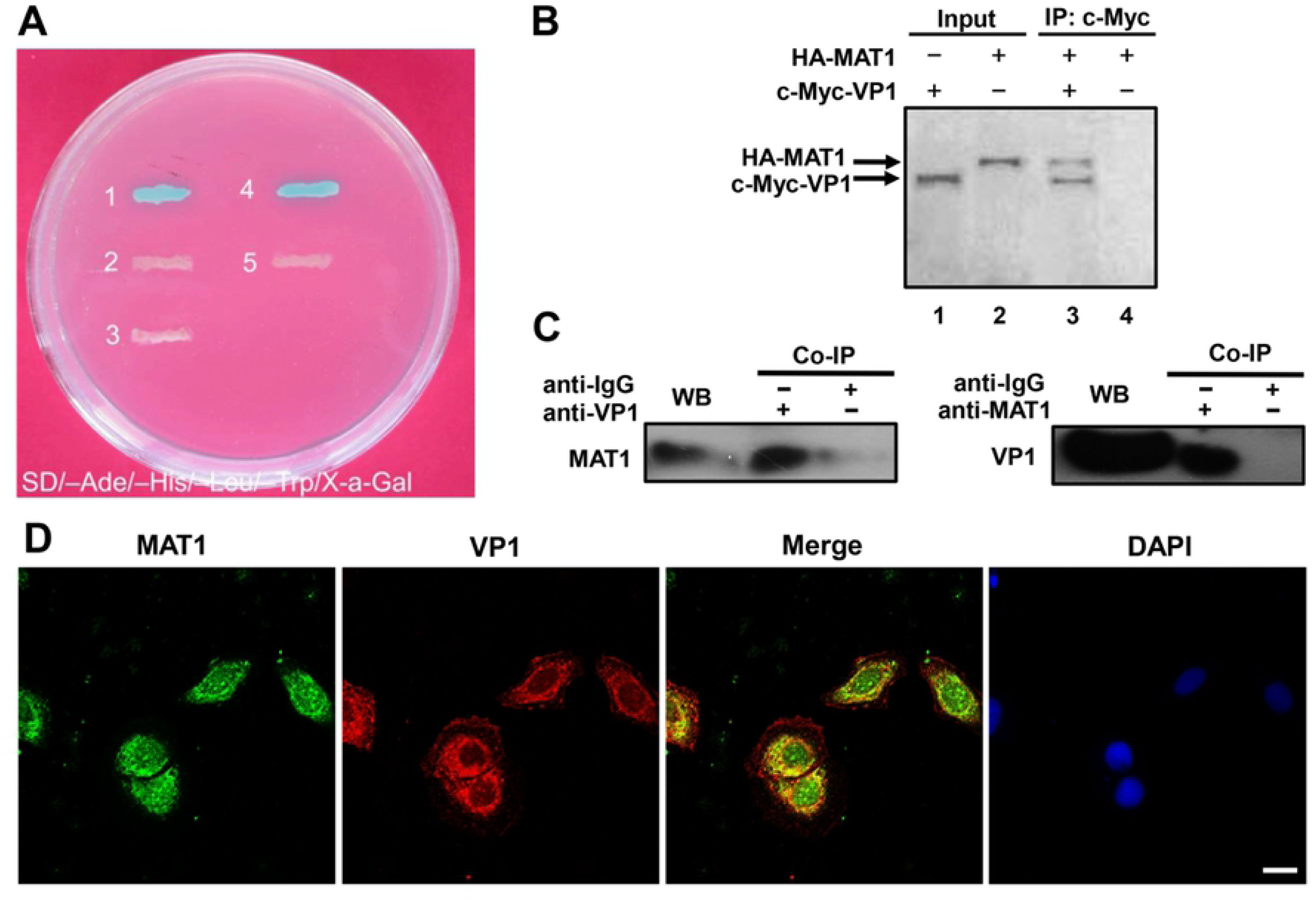
CVB3 VP1 may directly interact with MAT1. **(A)** Plasmids pGBKT7-VP1 and pGADT7-MAT1 were co-transformed with yeast strain AH109, and the positive result selected on high-stringency medium (SD/-Ade/-His/-Leu/-Trp). Interaction is reflected by blue color, while white colonies suggest no interaction. **(B)** the VP1-MAT1 interaction identified by transcription-translation system *in vitro*. Plasmids HA-MAT1 and c-Myc-VP1 were labeled with ^35^S-Methionine, and immunoprecipitated with antibody against c-Myc; the binding proteins with ^35^S-Methionine were analyzed by 10% SDS-PAGE and autoradiography. **(C)** Co-immunoprecipitation detected the interaction of VP1 and MAT1 in CVB3 infected cells. At 3 hours post infection, the cells lysates of mock and CVB3 infected cells were incubated with or without monoclonal antibody against MAT1 or irrelevant IgG affinity gel, and analyzed by western blotting with the specific antibodies after immunoprecipitation. **(D)** MAT1 and VP1 intracellular localization in CVB3 infected HPAC cells. Representative confocal immunofluorescence microscopic images of MAT1 and VP1 stained with rabbit anti-MAT1 (green) and mouse anti-VP1 antibodies (red), respectively, the MAT1 and VP1 images were also merged; the nuclei are labeled with DAPI. Scale bar = 10 µm. Values are shown as the mean ± SEM of three independent experiments. (\**P*<0.05, \*\**P*<0.01).

**Figure 3.**
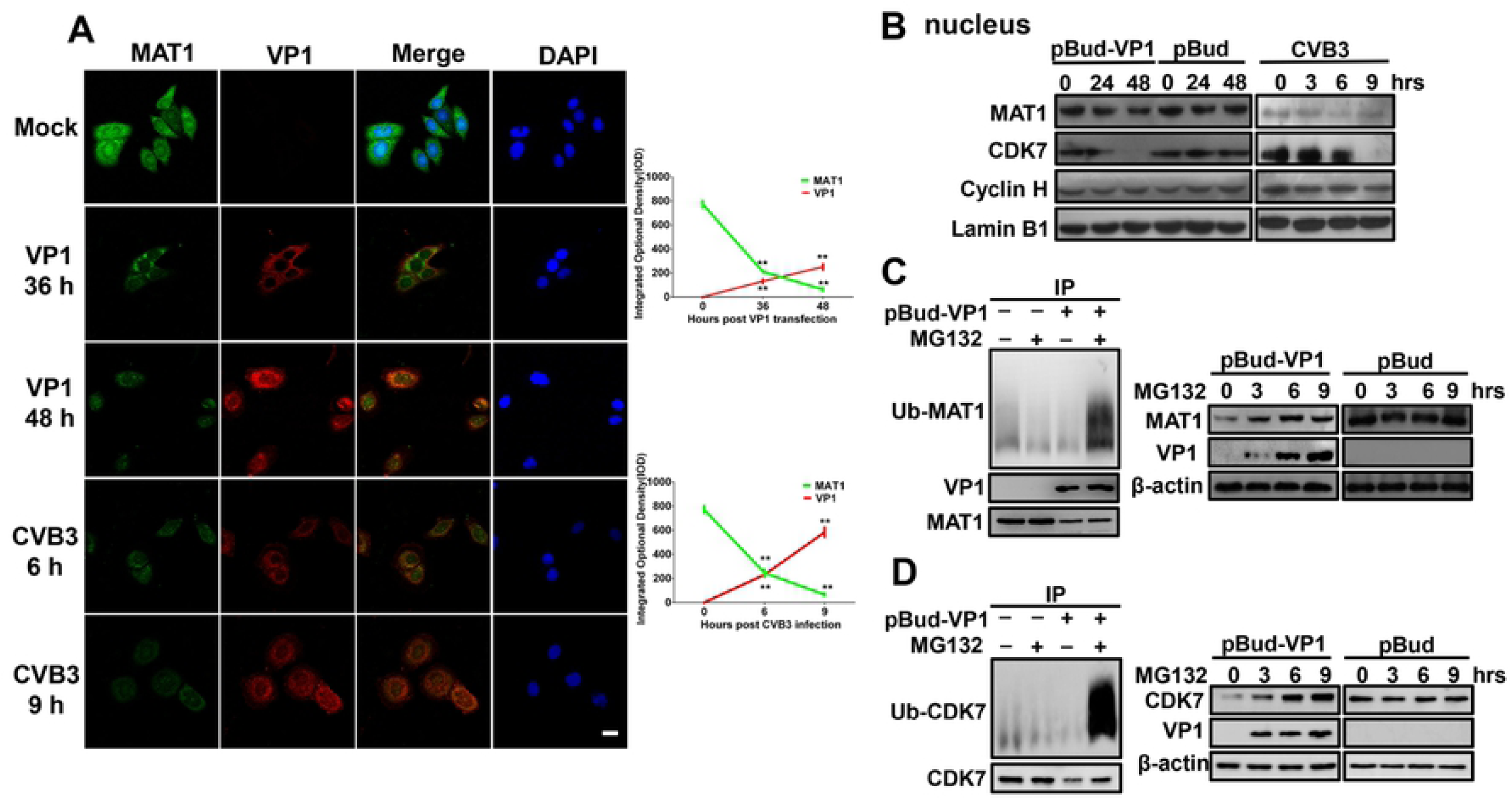
CVB3 VP1 impairs the assembly of a functional CAK complex. **(A)** Confocal microscopy analysis of the abundance of VP1 and MAT1 in HPAC cells, Cells were detected with monoclonal antibodies to MAT1 (Alexa-488) and polyclonal anti-VP1 (fluorescein; red), and counterstained with DAPI to show the nucleus. The MAT1 and VP1 images were merged. Scale bar: 10 µm. **(B)** Immunoblot analysis of the nuclear-localized accumulation of MAT1, CDK7, Cyclin H in CVB3 infected, VP1 and pBud transfected HPAC cells by specific antibodies. **(C)** CVB3 VP1 induces ubiquitination-proteolysis of MAT1. The cell lysates of pBud-VP1 and pBud transfected cells were incubated with monoclonal antibody against MAT1. The generated blot was then analyzed by immunoblotting with anti-ubiquitin antibody. **(D)** CVB3 VP1 induces ubiquitination-proteolysis of CDK7. Values are shown as the mean ± SEM of three independent experiments. (\**P*<0.05, \*\**P*<0.01).

We then targeted our investigation to the effects of CVB3 VP1 on cell cycle. To accomplish this, we first synchronized HPAC cell cultures in G1 phase, exposed them to VP1, and performed PI staining for analysis by flow cytometry. The resultant cell cycle profiles showed that in the VP1 transfection group, HPAC cells were significantly blocked at the G1 phase compared with the pBud group, while synchronized healthy cells continued their cell cycle after thymidine removal (Fig. 1D). These results aligned with experiments showing that CVB3 infection blocked cell cycle in both HPAC and HeLa cells (Supplemental Fig. 4). Moreover, PI analysis confirmed that the cell cycle was unaffected by exposure to CVB3 VP2, VP3, or VP4 (Supplemental Fig. 5). Taken together, these results revealed that VP1 inhibits HPAC and HeLa cell proliferation, exerts a strong cytopathic effect, and arrests cell cycle at G1/S phase.

**Figure 4.**
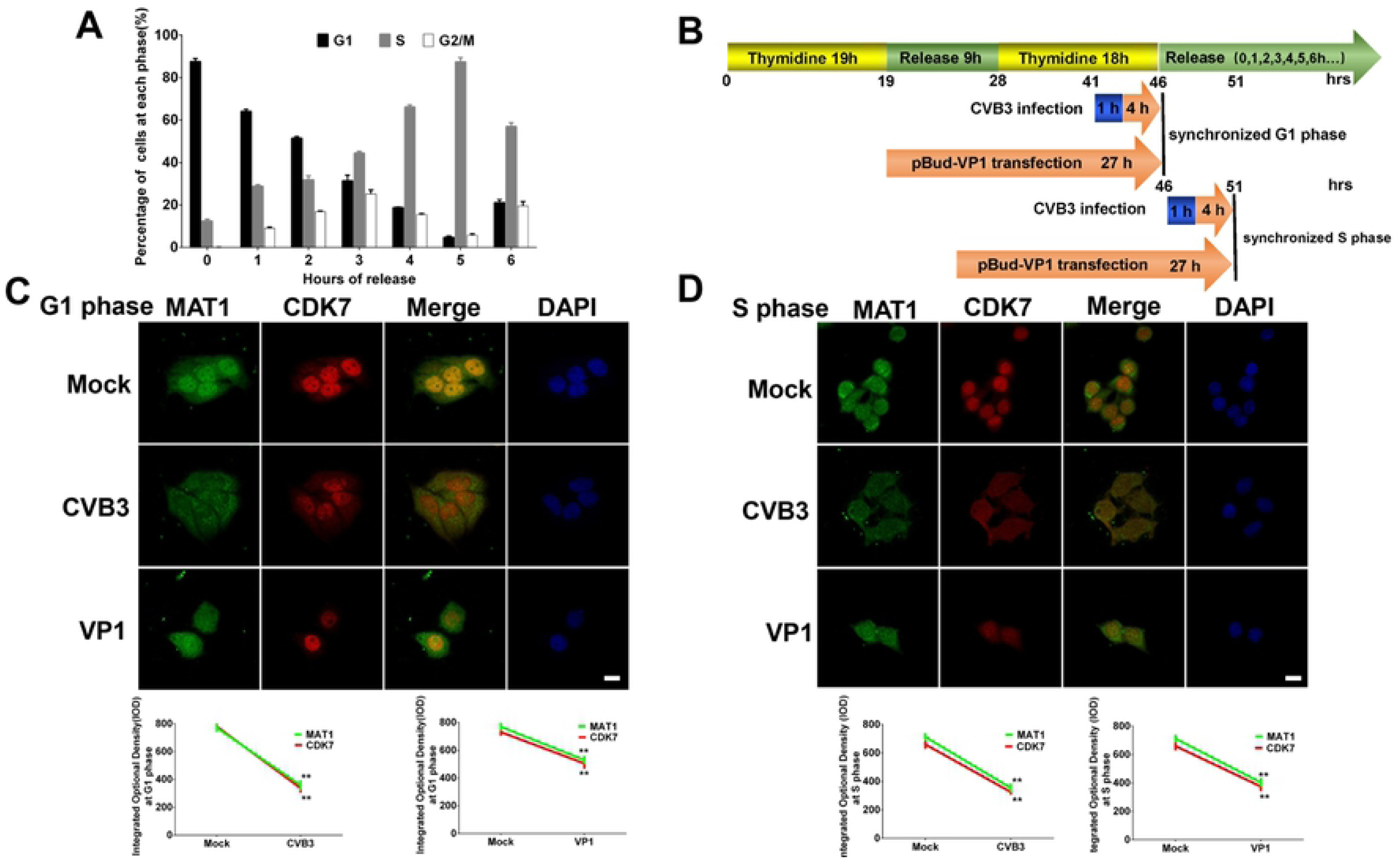
CVB3 VP1 alters the subcellular colocalization of MAT1 and CDK7 in G1/S phase. **(A)** Probe showing the time points when the cells were mainly at G1 and S phases by flow cytometry. The cells were synchronized using double thymidine block treatment, and released every hour. **(B)** Patterns of infection and transfection time points. The cells are synchronized at G1 (46 hrs) and S phase (51 hrs) by thymidine double blockade. **(C and D)** The alteration in spatial localization of MAT1 and CDK7 by thymidine double blockade at G1 and S phases, respectively. The cells were transfected with pBud-VP1 and infected with CVB3, and then imaged using confocal microscopy. Confocal image shows MAT1 (green), CDK7 (red), DAPI (blue) and merged views. Scale bar: 10 µm. The integrated optical density (IOD) was used to represent the fluorescence using Image Pro Plus (IPP). Values are shown as the mean ± SEM of three independent experiments. (\**P*<0.05, \*\**P*<0.01)

**Figure 5.**
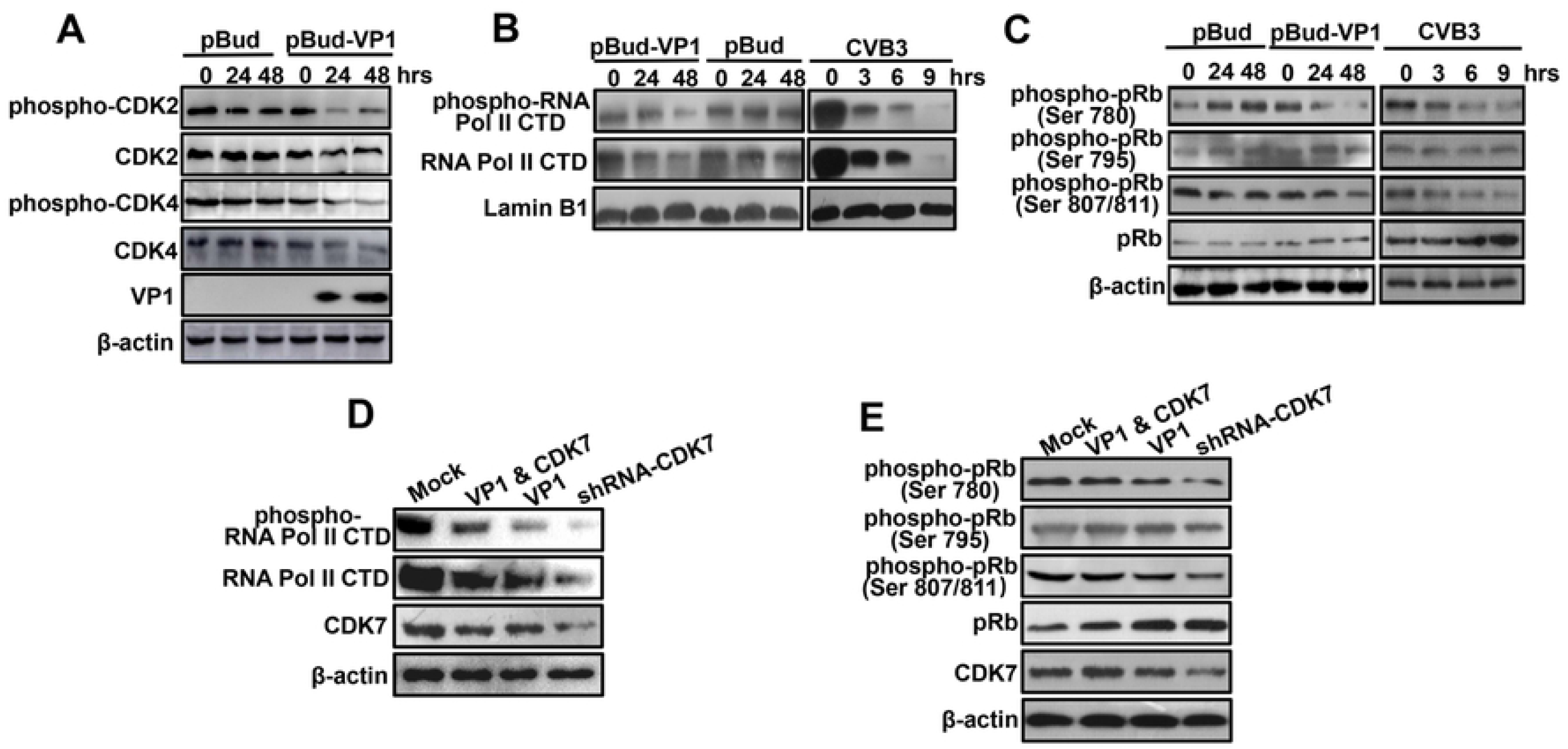
CVB3 VP1 could suppress CAK activity *in vivo*. **(A)** Time-course analysis of the accumulation of phosphorylated/nonphosphorylated CDK2 and CDK4 by indicated antibodies. **(B)** The expression level and phosphorylation of RNA Pol II CTD were detected by antibodies against RNA Pol II CTD and phospho-RNA Pol II CTD. CVB3 infection was the positive control. **(C)** Immunoblot analysis of the accumulation of phosphorylated/nonphosphorylated pRb with phospho-pRb Ser ^780, 795, 807/811^ antibodies. CVB3 infection was the positive control. **(D)** The abundance of RNA Pol II CTD, phospho-RNA Pol II CTD and CDK7 were analyzed by Western blot. **(H)** Western blotting analysis of the expression of phosphorylated/non-phosphorylated pRb and CDK7 with phospho-pRb Ser ^780, 795, 807/811^ and CDK7 antibodies.

### MAT1 was identified as a novel CVB3 VP1-binding protein

To obtain deeper insight into the potential mechanisms underlying CVB3 VP1 blockage of cell cycle at the G1/S phase and inhibition of cell proliferation, we screened for potential interacting proteins via yeast two-hybrid system, and subsequently identified a host cellular protein, MAT1, that acts as a VP1 binding partner (Fig. 2A). We then verified the two-hybrid binding of MAT1 to VP1, and autoradiography data confirmed that VP1 directly bound to MAT1 *in vitro* (Fig. 2B). Co-immunoprecipitation assays also confirmed the interaction in HeLa cells *in vivo*. Specifically, anti-VP1 was able to effectively precipitate MAT1, while anti-MAT1 also precipitated VP1, indicating that VP1 interacted with MAT1 *in vivo* (Fig. 2C). In light of these results, we then examined the cellular distribution and co-localization of VP1 and MAT1 by confocal microscopy. We observed that VP1 and MAT1 mainly shared the same spatial localization in the cytoplasm of the CVB3-infected HPAC cells (Fig. 2D) as well as in HeLa cells (Supplemental figure 6). Taken together, these results indicate that MAT1 is a binding partner for CVB3 VP1 both *in vitro* and *in vivo*.

**Figure 6.**
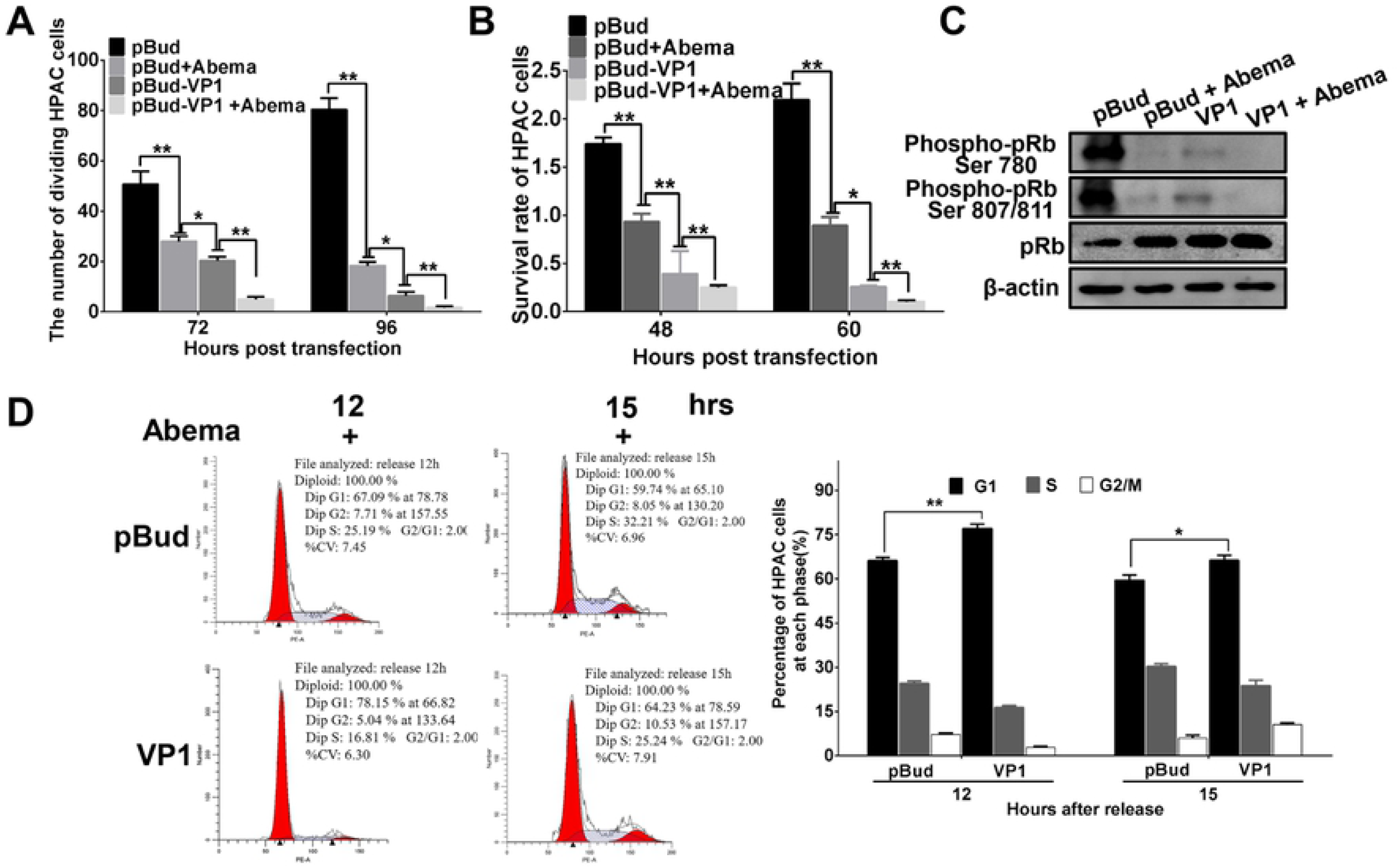
CVB3 VP1 inhibited the activity of CDK4/6 and Rb phosphorylation of the CDK-Rb signaling pathway in the G1/S transition. Abemaciclib (5 mg/ml stock solution in DMSO) was diluted to a final concentration of 10 µM in culture medium to treat cells. **(A)** Abemaciclib (Abema) further inhibits cell proliferation with VP1. An EdU assay was used to analyze the cell proliferation of Mock and pBud-VP1 transfected HPAC cells with or without abemaciclib treatment. **(B)** Abemaciclib further inhibits cell viability with VP1. A CCK-8 assay was used to analyze the cell viability of Mock and pBud-VP1 transfected HPAC cells with or without abemaciclib treatment. **(C)** Abemaciclib further inhibits phosphorylated pRb with VP1. Western blotting analysis of the expression of phosphorylated/nonphosphorylated pRb with phosphor-pRb Ser ^780, 807/811^ antibodies. CDK4/6 inhibitor further inhibits the cell cycle based on pBud-VP1. HPAC cells were treated with pBud and pBud-VP1 transfection within the treatment of the double-thymidine and abemaciclib, then released and separately collected at 12 and 15 hpi; the right figure is flow cytometric results. Values are shown as the mean ± SEM of three independent experiments. (\**P*<0.05, \*\**P*<0.01)

### VP1 induces CAK subunit ubiquitination to impair functional complex assembly

Given our findings of VP1 binding to MAT1, we then sought to determine the effects of this interaction on MAT1 function by observing the effects on MAT1 function and interactions. Confocal microscopy showed that VP1 expression increased in a time-dependent fashion with virus infection and pBud-VP1 transfection. Interestingly, with increasing expression of VP1, the expression of MAT1 decreased and even disappeared in VP1-transfected and CVB3-infected HPAC cells. We found a significant decrease in MAT1 signal intensity inversely related to an increase in VP1 intensity, thereby showing that VP1 decreased the cytoplasmic accumulation and nuclear localization of MAT1 (Fig. 3A). The effect of VP1 on MAT1 can also be observed in HeLa cells using immunohistochemical fluorescence staining (Supplemental figure 7). Furthermore, we used Western blots to assess the effects of VP1 on expression of CAK complex subunits MAT1, CDK7, and Cyclin H in the nucleus and cytoplasm in cells. The results indicated that nuclear expression of MAT1 and CDK7 declined with increasing time of VP1 transfection and CVB3 infection. Interestingly, we observed no detectable changes in the expression levels of Cyclin H in both group (Fig. 3B). In addition, MAT1 expression decreased and CDK7 / Cyclin H expression were undetectable in the cytoplasm (Supplemental Fig. 8). This result confirmed that MAT1, CDK7, and Cyclin H are all found in the cell nucleus, and that interference by VP1 attenuated the MAT1 expression in both the nucleus and cytoplasm, as well as the expression levels of nuclear CDK7.

**Figure 7.**
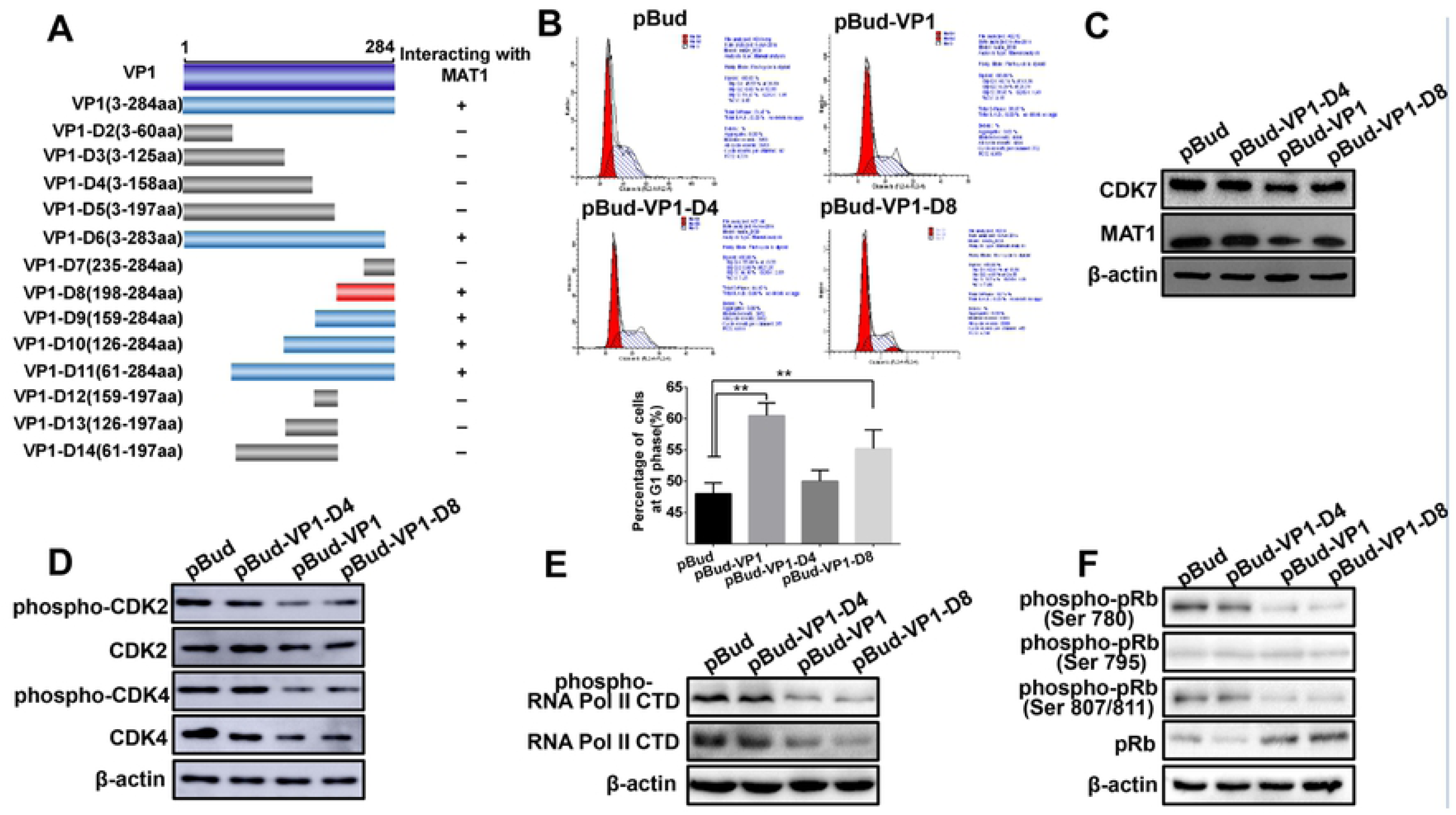
Mapping the minimum domain of VP1 interaction with MAT1. **(A)** Construction of a series of VP1 truncation/deletion mutants. The dark blue box is the full-length sequence encoded by VP1, the gray boxes are the VP1 deletion mutants that cannot interact with MAT1, the light blue boxes are the VP1 mutants that were detected to interact with MAT1, the D8 (red box) was the minimum domain of VP1 for the interaction (768-859 aa) in the VP1 C-terminus. **(B)** The interaction between VP1-D8 and MAT1 arrested cell cycle at the G1/S phase. pBud, pBud-VP1, pBud-VP1-D4 and pBud-VP1-D8 transfected cells for 48 h, and the cells were harvested and analyzed by flow cytometry. **(C)** VP1-D8 transfection down-regulated the expression of MAT1. Immunoblot analysis of the abundance of MAT1 in pBud, VP1, VP1-D4 and VP1-D8 transfected cells. **(D)** VP1-D8 transfection down-regulated the phosphorylation of CDK2 and CDK4. Western blot analysis of the accumulation of phosphorylated/nonphosphorylated CDK2 and CDK4 in different groups. **(E)** The accumulation of Pol II CTD, phospho-RNA Pol II CTD and β-actin were analyzed by Western blot. **(F)** Western blotting analysis of the expression of phosphorylated/nonphosphorylated pRb with specific antibodies. as in (D) using indicated antibodies. Values are shown as the mean ± SEM of three independent experiments. (\**P*<0.05, \*\**P*<0.01).

**Figure 8.**
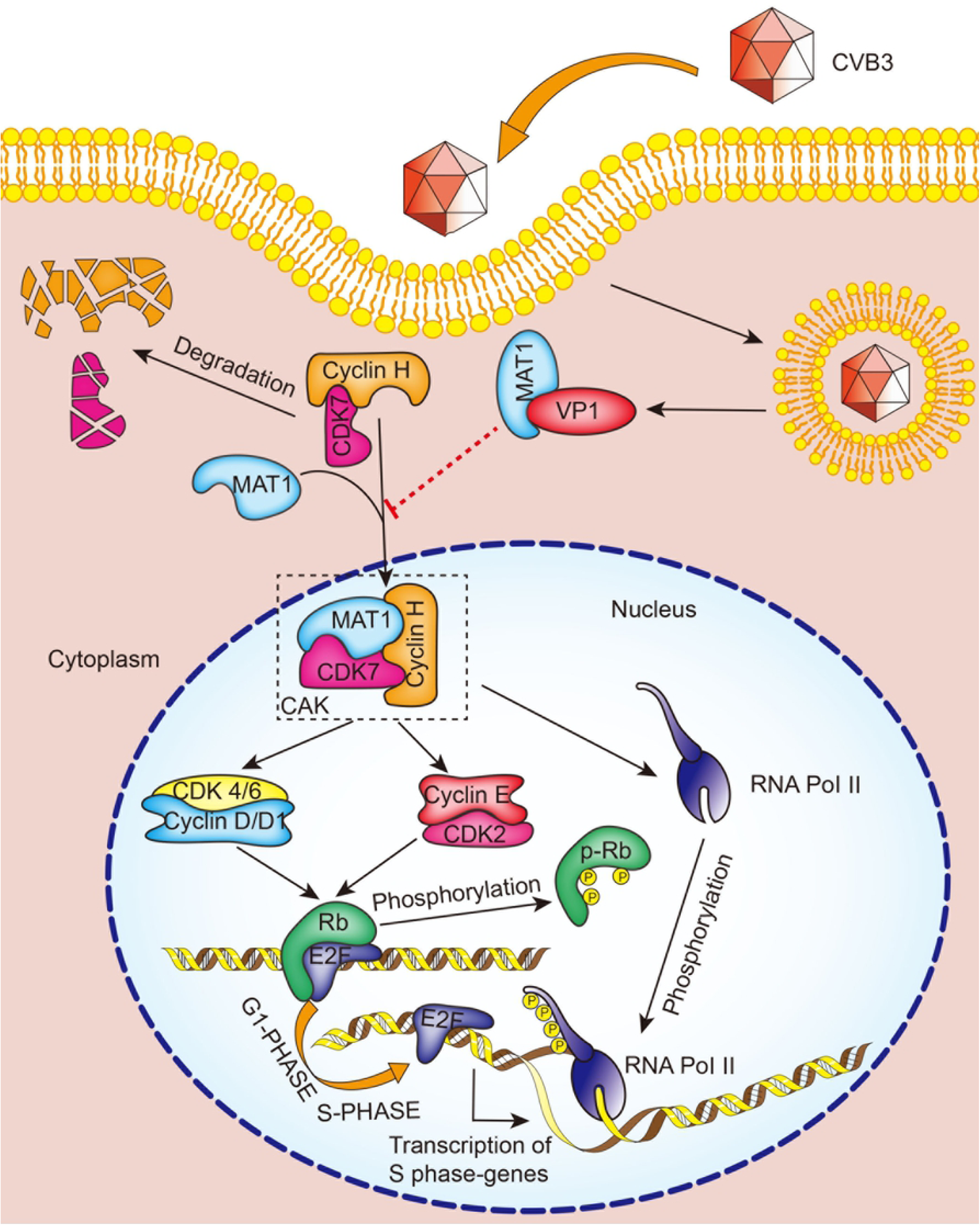
A schematic diagram representing how CVB3 VP1 inhibits cell proliferation via impairing the CAK complex through its interaction with MAT1. As CVB3 invades HPAC and Hela cells, CVB3 VP1 interacts competitively with MAT1, leading to the degradation of CDK7, thus the assembly of the CAK complex is inhibited, and the activity of CAK is simultaneously inhibited. CAK affects the activity of cyclin-dependent kinases CDK2 and CDK4, and then inhibits the phosphorylation level of pocket Rb protein, inducing the persistent binding of Rb and E2F protein, and finally inhibits the expression of downstream transcription factors by E2F and cell cycle arrest occurs in the G1 phase. CAK also affects the phosphorylation of its substrate RNA Pol II CTD Ser ^5^, which eventually leads to the inability of RNA Pol II to participate in cell transcription and cell cycle arrest at the G1 phase.

To further explore the reasons why expression of CAK complex proteins MAT1 and CDK7 were down-regulated by VP1, we conducted ubiquitin proteasomal degradation assays. We found that in VP1-transfected cells, anti-MAT1-immunoprecipitated MAT1 was ubiquitinated and appeared as a smear of degraded protein (Fig. 3C left). Concurrently, the MAT1 band intensity substantially increased by the addition of MG132 proteasomal inhibitor, and the intensity of MAT1 ubiquitination also progressively increased with prolonged incubation with MG132 (Fig. 3C right). As shown in Fig. 3D, VP1 exposure correspondingly led to CDK7 degradation, as observed through ubiquitin proteasomal degradation assays (Fig. 3D). Taken together, these results strongly suggest that CAK complex assembly and function is likely impaired through the ubiquitination and proteasomal degradation of its subunits during exposure to VP1.

### CVB3 VP1 interferes with subcellular colocalization of MAT1 and CDK7

In order to determine whether VP1 impaired MAT1 interactions with other subunits, we observed colocalization of MAT1 and CDK7 in CVB3 VP1-transfected cells in G1 and S phase of the cell cycle. To this end, we first determined the optimal release time point from G1 to S phase after thymidine double blocking, thus synchronizing cells in the G1 and S phase transition. The results showed that the highest percentage of synchronized healthy cells in the G1 and S phase could be observed at 0 h and 5 h after release, respectively (Fig. 4A). We next transfected VP1 and infected CVB3 (Fig. 4B), and observed through confocal microscopy that in the G1 phase, VP1 and CVB3 induced the transfer of a large proportion of nuclear-localized MAT1 to the cytoplasm, concurrent with decreased accumulation of MAT1 in both the nucleus and cytoplasm and decreased nuclear levels of CDK7 (Fig. 4C). In contrast, among cells synchronized in the S phase, VP1 and CVB3 attenuated nuclear and cytoplasmic MAT1 accumulation but did not result in MAT1 export from the nucleus to the cytoplasm. Interestingly, VP1 and CVB3 induced a partial transfer of nuclear-localized CDK7 to the cytoplasm (Fig. 4D). Based on these findings, we therefore concluded that the interaction between CVB3 VP1 and MAT1 not only reduced the levels of MAT1 and CDK7, but also substantially altered the subcellular colocalization patterns of MAT1 and CDK7, likely disrupting the function of CAK complex.

### CVB3 VP1-MAT1 interaction suppresses CAK activity via CDK7 *in vitro*

Since the results above suggested that the VP1-MAT1 interaction disrupted the assembly of the CAK complex, we addressed whether the interaction between VP1 and MAT1 also reduced CAK activity in VP1-transfected or CVB3-infected cells. CAK indirectly phosphorylates Rb via Cyclin-dependent kinases (CDKs), and the activation of CDK relies on T-loop phosphorylation by CAK[15, 35]. We therefore assessed the expression and phosphorylation levels of CDK2 or CDK4. The results showed that the protein expression and phosphorylated protein levels of CDK2 or CDK4 were progressively attenuated over the 48 h time course in pBud-VP1 transfected cells (Fig. 5A).

Previous studies have reported that the carboxy-terminal domain (CTD) of the largest subunit of RNA Pol II is a canonical direct substrate of the CAK complex, and that the phosphorylation of RNA Pol II Ser^5^ and Ser^7^ residues is primarily mediated by CDK7 activity [19, 36, 37]. Therefore, we evaluated CAK activity in cells exposed to VP1 through quantification of RNA Pol II CTD Ser^5^ phosphorylation levels. After nucleoplasm separation, we found that RNA Pol II CTD and phosphorylated RNA Pol II CTD were both down-regulated in the nucleus (Fig. 5B). In addition, we detected the phosphorylation levels of the CAK complex indirect substrate, pRb, which is the substrate of Cyclin D/CDK4 or Cyclin E/CDK2, both of which are direct substrates of the CAK complex[38]. These results showed that both pRb Ser^780^ and pRb Ser^807/811^ phosphorylation were down-regulated in the pBud-VP1 transfected or CVB3 infection groups. In contrast, pRb total protein levels were significantly elevated in cells exposed to VP1. However, there was no significant change in pRb Ser^795^ site phosphorylation among any of the groups (Fig. 5C).

To verify whether CDK7 mediated the phosphorylation of RNA Pol II CTD and pRb in VP1-transfected cells, we silenced the CDK7 gene in HPAC cells using an shRNA vector. We also overexpressed CDK7 in HPAC cells to verify whether increased CDK7 activity can rescue CAK function. The results showed that phosphorylation of RNA Pol II CTD was further decreased in cells co-transfected with the CDK7 silencing plasmid and VP1, compared with that of cells transfected with VP1 alone. In agreement with these results, phosphorylation of RNA Pol II CTD was recovered in cells co-transfected with VP1 and the CDK7 over-expression plasmid (Fig. 5D). In addition, lower phosphorylation levels of pRb Ser^780^ and pRb Ser^807/811^ were observed in co-transfections of VP1 and the CDK7 silencing construct compared to those transfected with VP1 alone. Similarly, pRb Ser^780^ and pRb Ser^807/811^ phosphorylation was apparently rescued to the high level observed in control cells after co-transfection with VP1 and CDK7 over-expression plasmid, while total pRb protein showed the opposite trend. Furthermore, we were unable to detect any changes in phosporylation of pRb at Ser^795^ (Fig. 5E). Collectively, these results indicate that the VP1-MAT1 interaction indeed impairs the catalytic activity of the CAK complex by interfering with phosphorylation of CDK2 or CDK4, as well as CDK7 kinase activity towards RNA PoI II and pRb phosphosites Ser^780^ and Ser^807/811^.

### CVB3 VP1 inhibits the activity of CDK4/6 and Rb phosphorylation of CDK-Rb signaling pathway proteins in the G1/S transition

To further investigate whether CVB3 VP1 perturbs the activity of CDK4/6 and the Rb phosphorylation of the CDK-Rb signaling pathway in the G1/S transition, we conducted a series of assays to compare the effects of the CDK4/6 inhibitor abemaciclib with that of VP1 on the inhibition of HPAC cell proliferation and CDK4/6 activity. The results showed that abemaciclib and CVB3 VP1 similarly reduced cell proliferation in the pBud group, while exposure to VP1 alone led to a more powerful inhibitory effect than pBud plus abemaciclib together (Fig. 6A and B). These results were also reflected by comparable patterns of attenuation to pRb phosphorylation, with greater inhibition associated with dual treatments compared to VP1 exposure alone (Fig. 6C). Moreover, among cells treated with abemaciclib, flow cytometry showed that a greater proportion of cells also transfected with VP1 were arrested at the G1 phase compared to cells treated only with abemaciclib at both 12 and 15 h after release from double thymidine (Fig. 6D). These results collectively show that VP1 indeed interrupts CDK4/6 activity and Rb phosphorylation of CDK-Rb pathway proteins, thereby leading to inhibition of cell proliferation.

### CVB3 VP1 C-terminal domain (198-284 aa) is required for interaction with MAT1

Having established the interaction between VP1 and MAT1, we then used yeast-two hybrid to identify which regions of VP1 are required for binding with MAT1. Co-transformation of a series of VP1 truncation/deletion variants with pGADT7-MAT1 revealed that the VP1 variant carrying amino acids 198-284 interacted with MAT1, while segments spanning amino acids 3-197 showed no interaction (Fig. 7A). Further, we examined the CVB3 VP1-D8 domain to determine whether it exhibited similar functions to that of full-length VP1. As shown in Fig. 7B, VP1-D8 functioned comparably to VP1 in the inhibition of cell cycle at the G1 phase, and similarly attenuated the accumulation of MAT1, whereas VP1-D4 did not (Fig. 7C). To strengthen the evidence for this function, we investigated whether VP1-D8 was also able to affect activity towards CAK complex substrates. For this purpose, we used Western blot assays to confirm that VP1-D8 bound to MAT1 similarly to the full length VP1, and that it also reduced phosphorylation of CDK2/4, RNA Poly II CTD, and pRb, while VP1-D4 and pBud did not (Fig. 7D, E, and F). Taken together, these data collectively indicate that VP1-D8 comprises the minimum VP1 domain necessary to block the cell cycle at G1 phase, attenuate CAK complex phosphorylation activity towards its substrates via MAT1 binding, and ultimately suppress cell proliferation.

## Discussion

In this study, we present the first report of which we are aware detailing the mechanisms of CVB3 structural protein VP1 inhibition of pancreatic cell proliferation. We discovered VP1 interacts with MAT1, leading to down-regulation of MAT1 and CDK7, and ultimately impairing CAK complex formation. Furthermore, we pinpointed the minimum functional domains of VP1 that engage in this interaction.

However, deregulation of cell proliferation leads to diseases characterized by either excess proliferation or cell loss, an effect that is induced by some viral infections due to exploitation of replication machinery to benefit for viral replication. Viral targeting of the cell cycle as a core regulatory component of cell proliferation has been widely studied among other viruses. For example, latent protein 3C (EBNA3C) of Epstein-Barr virus (EBV), the first reported human tumor virus, can directly bind to CDK2, and also cooperate with master transcription factor Bcl6 to regulate the expression of CDK2, thereby promoting cell proliferation [39]. In contrast, Hepatitis C virus (HCV) reduces CD 8^+^ T cell proliferation, and leads to dysfunction in HCV chronic infection [40]. Furthermore, HCV decreases the proportion of infected cells in G1 and S phase commensurate with accumulation of G2/M cells [41]. Indeed, most viral proteins reported to participate in regulating cell proliferation are non-structural, whereas viral structural proteins more commonly contribute functions such as protection of the viral genome against inactivation by nucleases, viral attachment to host cells, providing a symmetry to virus particle structure, which can also provide crucial antigenic characteristics. However, it remains largely unknown whether and how structural proteins can interact with host proteins to regulate host cell functions, such as cell cycle regulation or stress response.

In HPAC and HeLa cell models, our results indicated that the CVB3 capsid protein VP1 induced cell cycle arrest, leading to inhibition of cell proliferation and cytopathic effects. Furthermore, this process is mediated by interaction with CAK complex assembly factor, MAT1. Our findings showed that structural viral proteins can inhibit cell proliferation via cell cycle arrest. A widely accepted mechanism of G1/S phase arrest is through inactivation of cyclinD-CDK4 phosphorylation, phosphorylation of pRb, and concomitant E2F activation[42, 43]. However, our study revealed that VP1 directly or indirectly inactivated phosphorylation of CDK2 and CDK4, pRb, and RNA Pol II to arrest the G1 phase, thus suggesting a new model for interference in cell proliferation.

We then investigated how the interaction of VP1 with MAT1 leads to the inhibition of cell cycle and found that MAT1 is expressed in the cytoplasm and nucleus in the G1 phase, but primarily localized to the nucleus. Interference by CVB3 VP1 decreased MAT1 expression, destabilized CDK7, and impaired CAK complex formation in the nucleus. While in the S phase, CDK7 is expressed only in the nucleus, but readily diffused into the cytoplasm due to interference of VP1, which aligned with our results showing that CDK7 binding to MAT1 decreased upon pBud-VP1 transfection. Among several known functions of MAT1, its C-terminal domain activates CDK7 phosphorylation and stabilizes Cyclin H / CDK7 binding to the TFIIH core through interaction with XPB and XPD [44]. Interestingly, we observed VP1 bind to the C-terminal of MAT1 (data not shown), thus indicating that interaction between VP1 and MAT1 can disrupt CAK complex assembly, and decrease phosphorylation activity by CDK7. Moreover, we found that CDK7 was degraded through the ubiquitination pathway in the presence of VP1.

In addition to preventing CAK complex assembly, VP1 binding to MAT1 could also impair CAK complex activity by interfering with CDK7-mediated phosphorylation. We found that phosphorylation of the direct or indirect CAK substrates CDK2, CDK4, and RNA pol CTD were all attenuated following transfection of VP1, and that inhibition of pRb phosphorylation involved its Ser^780^ and Ser^807/811^ sites. Overexpression or silencing of CDK7 revealed that VP1 / MAT1 interactions led to inhibition of CAK activity via CDK7 degradation. Furthermore, we found that VP1-D8 was the minimally required domain for VP1-mediated blockade of the cell cycle at G1/S phase, to successfully suppress cell proliferation and inhibit CAK complex activity.

We subsequently determined that CVB3 VP1 also inhibited CDK4/6 activity and Rb phosphorylation in the G1/S transition. In fact, the pocket protein Rb is among the most functionally essential proteins phosphorylated by CDK4/6 and CDK2 to regulate cell cycle[45]. Given that VP1 inhibition of CDK4/6 activity can attenuate pRb phosphorylation, thereby leading to arrest cell cycle in the G1 phase, we thus conclude that CVB3 VP1 can potentially inhibit cell proliferation through a MAT1-mediated CAK-CDK4/6-Rb signaling pathway.

Given previous studies showing that CVB3 can cause acute pancreatitis in humans and mice, and that TNFα and IL6 are implicated in the development of this disease [64, 65], it is likely that cytokines are also involved in virus-induced inhibition of cell proliferation. Moreover, the expression of IL10 may suppress acute pancreatitis [66], and notably, *Sawada* found that IL-10 and its downstream STAT3 pathway regulate the proliferation of cells infected with HTLV-1 [46]. In addition, IL-6-deficient mice infected with influenza virus were found to produce high levels of TGF and enhanced the proliferation of lung fibroblasts [47]. In contrast, we found that in the G1 phase of VP1 transfection, inflammatory cytokines (TNFα and IL-6) were up-regulated, while anti-inflammatory cytokine (IL-10) was down-regulated (Data not shown). Taken together with our previous work showing that CVB3 infection caused pancreatitis, we thus hypothesized that the changes in the accumulation of these inflammatory factors induced by VP1 transfection in HPAC cells synergistically blocked the cell cycle in the G1 phase, potentially resulting in pancreatic inflammation. However, we cannot yet exclude the possibility of additional contributing factors, and ongoing research will clarify the full extent of host proteins participating in CVB3 induction of pancreatitis.

In this study, we provide evidence that direct interaction between CVB3 VP1 and MAT1 produces an inhibitory effect on HPAC cell proliferation by blocking the MAT1-mediated CAK-CDK4/6-Rb signaling pathway required for CAK complex assembly and activity, which thereby results in cell cycle arrest during the G1 phase in pancreatic cells.

## Material and Methods

### Cell culture, synchronization and virus

The human pancreatic adenocarcinoma cell line HPAC, HeLa and 293A cells were obtained from the American Type Culture Collection (Manassas, VA, USA). Cells were grown in a 5% CO2 incubator at 37°C, and maintained in Dulbecco’s modified Eagle’s medium (BI, Israel) and DF12 (BI, Israel) supplemented with 10% heat-inactivated 10% fetal bovine serum (BI, Israel).

HPAC and HeLa cells were synchronized at the G1 phase using double thymidine block treatment. Briefly, cells in six-well plates grew to 40% confluency, then thymidine was added to a 2 mM final concentration and cells were incubated for 34 h (HPAC) or 19 h (HeLa). Subsequently, the medium was aspirated with thymidine and the cells were washed twice with PBS, after releasing for 10-12 h (HPAC) or 9 h (HeLa). In the second thymidine block, thymidine was added to the cells for a final concentration of 2 mM and incubated for 33 h (HPAC) or 18 h (HeLa).

CVB3 (Nancy strain; GI:323432) was propagated in HeLa cells and purified using a method previously described by Henke et al. [48]. Cells were infected with CVB3 (MOI of 5) throughout the study.

### Plasmids construction

To construct yeast two-hybrid plasmids, VP1 and MAT1 genes were cloned into Vector pGBKT7 (Clontech, USA). We generated pGBKT7-VP1 and pGADT7-MAT1 plasmids to map the regions in VP1 that were required for their interaction in the yeast-two hybrid system. Briefly, the different truncation/deletion mutants of CVB3 VP1 were cloned into pGBKT7 vector (between the Nde I & Pst I sites) in frame with the vector’s GAL4 DNA Binding Domain (BD), and MAT1 was expressed in the pGADT7 vector (between the EcoR I & BamH I sites) in frame with the vector’s GAL4 activation domain (AD). The deletion mutants of VP1 were separately generated by PCR with pGBKT7-VP1 as the template. VP1, VP1-D2, VP1-D3, VP1-D4, VP1-D5, VP1-D6, VP1-D7, VP1-D8, VP1-D9, VP1-D10, VP1-D11, VP1-D12, VP1-D13, VP1-D14 were separately cloned into pGBKT7 vector. VP1, VP1-D4 and VP1-D8 were cloned into pBudCE4.1 (pBud) vector. CDK7 was cloned into pCAG-Flag and VP2, VP3, and VP4 were cloned into pAdtrack-Flag. All cDNAs were PCR-amplified using Phanta Max Super-Fidelity DNA Polymerase (Vazyme, China). The PCR fragments were ligated to the expression plasmid using the ClonExpress II One Step Cloning Kit (Vazyme, China). All primers in this study used for plasmid construction are listed in Supplemental Table 1.

### Transient transfection

The plasmids were transfected into HPAC and HeLa cells according to the manufacturer’s instructions (Thermo Fisher Scientific, USA). Cell (2×10^5^) were seeded into six-well plates, and the fusion degree of cell monolayers reached 70%-80% after 24-hour incubation. Then, 400 µl DMEM or DF12, 4 µg plasmid and 6 µl Turbofect (Thermo Fisher Scientific, USA) were combined. The DNA-Turbofect mixture was allowed to sit for 15-20 minutes, then the cells were washed twice with PBS, and 4 ml DMEM or DF12 supplemented with 10% fetal calf serum and DNA-TurboFect mixture were added.

### Recombinant adenovirus construction, production and use

To generate the adenovirus to express *VP2, VP3* and *VP4* proteins, VP2, VP3 and VP4 genes from CVB3 were cloned into the pAdtrack-CMV vector. The newly constructed pAdtrack-*VP2/VP3/VP4* vectors were linearized with Pme I digestion and then cotransformed with pAdEasy-1 vector (AdEasy Adenoviral vector system; Stratagene, USA)[49] into *E. coli* BJ5183 for homologous recombination, creating *VP2/VP3/VP4* Ad vectors. Additionally, all recombinant adenoviruses were packaged and propagated using 293A cells. The infection was performed as described in previous studies[50, 51]. The titer of viral stocks was assessed using real-time PCR.

### Cell proliferation assay

The effect of cell proliferation was analyzed using a CCK-8 assay. HPAC and HeLa cells were seeded in 96-well plates at the density of 3,000 cells into each well, and after nearly 24 hours of incubation, the two cells were transfected with pBud or pBud-VP1 plasmids at different times. A total of 10 µl CCK-8 solution (Vazyme, China) was added each well in the plate and incubated for 2-4 hours in a 5% CO2 incubator at 37°C, and the absorbance was recorded at 450 nm in a microplate reader (SpectraMAX^®^ Paradigm^®^).

### EdU incorporation assay

The capacity of DNA replication was determined using a 5-Ethynyl-2’-deoxyuridine (EdU) assay according to the manufacturer’s protocol (RiboBio, China). Cells in a logarithmic state were seeded in a 96-well culture plate at the density of 5,000 cells per well and incubated for 24 h, then transfected with plasmids pBud and pBud-VP1 for 0, 24, 48, 72 h. Next, cells were treated with 50 µM EdU for 4 h (HPAC) or 2 h (HeLa), then fixed with 4% polyformaldehyde for 30 min, and finally incubated with 2 mg/ml Glycine for 5 min. After treating with 0.5% Triton X-100 for 10 min, cells were stained with Apollo 567 for 30 min, and then cell nuclei were incubated with Hoechst33342 for 30 min. All samples were measured using a fluorescence microscope (Olympus IX17).

### Flow cytometry

HPAC cells (2×10^5^) were seeded in each well of a six-well plate, then treated with plasmid (pBud and pBud-VP1) transfection and CVB3 infection within the treatment of double-thymidine. Cells were then released and collected at indicated time, the number of collected cells ranged from 1-5×10^6^. The collected cells were washed with cold PBS, fixed using 75% ethanol and stored overnight at 4°C. Before detection, the fixed cells were washed with cold PBS, bathed with RNase A at 37°C, and the cell adhesives were filtered with 400 mesh screen, then stained with propidium iodide (PI) for analysis, and kept away from light for 30 minutes. Flow cytometry analysis was used to determine cell percentages at different stages of the cell cycle (BD FACSCVerse).

### Antibodies

Proteins were detected using the following primary antibodies: anti-enterovirus VP1 clone 5-D8/1 antibody purchased from Dako (Denmark); Mouse anti-Ub, Mouse anti-MAT1, Mouse anti-CDK7, and Mouse anti-Cyclin H antibodies purchased from Santa Cruz (USA); Mouse monoclonal to RNA polymerase II CTD and Rabbit monoclonal to RNA polymerase II CTD (phosphor S5) antibodies purchased from Abcam (USA); Mouse anti β-actin purchased from Proteintech; Rabbit monoclonal CDK2 and CDK4, Rabbit monoclonal phosphor-CDK2 and CDK4, Mouse monoclonal pRb, Rabbit monoclonal pRb-phospho Ser780, pRb-phospho Ser795 and pRb-phospho Ser807/811 antibodies purchased from Cell Signaling (USA); Rabbit polyclonal Lamin B1, Mouse monoclonal c-Myc, Mouse monoclonal HA-Tag and Mouse Monoclonal Flag antibodies purchased from Sigma (USA); Goat anti-mouse IgG horseradish peroxidase-conjugated secondary antibody and goat anti-rabbit IgG horseradish peroxidase-conjugated secondary antibody purchased from Thermo Fisher Scientific (USA).

### Western blot

The total protein was washed with PBS and lysed with lysis buffer (150 mM NaCl, 20 mM Tris HCl, 0.1% NP-40, pH 7.4). The protein supernatant was collected after centrifugation. The same amount of protein was injected into a 10% SDS-PAGE gel for separation and transferred to a polyvinylidene fluoride (PVDF) membrane (Millipore, Germany). Cell membranes were blocked at room temperature for 2 hours with 5% skim milk and incubated with primary antibodies overnight at 4°C, then incubated with secondary antibodies for 1-2 hours at room temperature. The protein was detected with an enhanced Chemiluminescence Kit (Thermo Fisher Scientific, USA).

### Co-immunoprecipitation

The VP1 interactome was captured from the cells with Mouse anti-enterovirus VP1. Briefly, CVB3-infected cells at 3 hpi were lysed with lysis buffer (150 mM NaCl, 20 mM Tris HCl, 0.1% NP-40, pH 7.4) and nuclei were removed by a 10 min low-speed centrifugation step. Precleared cell lysates were incubated with or without monoclonal antibodies against MAT1 (1:500) or irrelevant IgG for 1 h, and for an additional hour with 20 µl Protein A/G Agarose beads (Thermo Fisher Scientific, USA) by gentle rotation at 4 °C. The beads were washed three times in lysis buffer with protease inhibitors, and two times in a wash buffer (100 mM Tris pH 7.4) with 500 mM LiCl before suspending in 5× SDS sample buffer. The supernatant and crude extracts were analyzed using 10% SDS/PAGE, transferred to PVDF membranes (Millipore, Germany) and detected with specific antibodies against VP1 (1:1,000).

### Ubiquitin proteasomal degradation assay

The ubiquitin proteasomal degradation was induced by HPAC cells with 10 µm MG132. Briefly, HPAC cells were seeded in six-well culture plates for 24 h and transfected with pBud-VP1 for 40 h, and then incubated with MG132 at a 10 µm final concentration for 8h. Cells were lysed with lysis buffer and supplemented with protease inhibitor PMSF, the cell lysate was centrifuged, and the supernatant was collected. Before incubation with 20 µl Protein A/G Agarose beads and specific antibodies, antibody against MAT1 or CDK7 and Protein A/G Agarose beads were additionally combined for 4 h, then incubated with protein supernatant overnight at 4°C. Subsequently, the protein supernatant was washed three times with ice-cold lysis buffer and eluted in 5× SDS sample buffer. Western blot was performed for anti-Ub and anti-β actin antibodies with protein samples.

### Indirect immunofluorescence labelling and confocal microscopy

To visualize the co-localization of VP1 and MAT1 in HPAC cells and the changes of their expression, the CVB3-infected HPAC cells and pBud VP1-transfected HPAC cells were incubated with fluorescein conjugated goat anti-rabbit IgG (1:200) and rhodamine-conjugated goat anti-mouse IgG (1:200) at room temperature for 1.5 h with primary antibodies. The primary antibodies used were rabbit anti-MAT1 (1:50), Mouse anti-enterovirus VP1 clone 5-D8/1 (1:50) and Mouse anti-CDK7 (1:50). The cells were rinsed extensively with PBS, and DAPI was used for counterstaining. The stained slides were analyzed with a FLUOVIEW FV1000 confocal laser scanning microscope (Olympus) with Olympus FV1000 software.

### Yeast two-hybrid screen and assay

Yeast two-hybrid screen and assay were performed as previously described[14]. In order to determine which region of CVB3 VP1 interacts with MAT1, various truncated deletion mutants of VP1 were cloned into pGBKT7 bait vectors with the Matchmaker System. The yeast strain AH109 was co-transformed with pGADT7-MAT1 prey vector and bait vectors for expression of different truncated coding sequences of VP1 using lithium acetate. Transformants positive for prey-bait interaction were grown on selection plates lacking tryptophan, leucine, adenine and histidine but containing X-α-Gal.

### Protein-protein binding assay *in vitro*

An *in vitro* protein-protein binding assay was performed as previously described[14]. pGBKT7-VP1 bait vector and pGADT7-MAT1 prey vector were used as templates to transcribe and translate *in vitro*, then labeled with ^35^S-Methionine (Amersham Pharmacia Biotech) *in vitro* transcription-translation system (Promega), respectively, to obtain ^35^S-labeled fusion proteins HA-MAT1 and c-Myc-VP1. The ^35^S-Methionine-labeled HA-MAT1 and c-Myc-VP1 were incubated at room temperature, then incubated with antibody against c-Myc (Clontech) in lysis buffer, subsequently mixed with protein A/G plus-agarose (Thermo Fisher, USA) and incubated for 3 h at 4°C. The beads were washed three times with lysis buffer. The radioactive antibody–protein complexes were eluted, and subjected to SDS/PAGE and then autoradiography.

## Statistical analysis

All statistical analyses were performed using GraphPad Prism 6.0 and SPSS software. The raw data are expressed as means ± standard error of the mean (s.e.m.) of at least three independent repeats. Unpaired two-tailed Student’s *t*-tests were applied to analyze the differences for all comparisons in SPSS version 17.0. Band density on western blot was evaluated and quantified using Image J software.

### Acknowledgements

This study was supported by the National Natural Science Foundation of China (31760261 and 31660035), the Science and Technology Research Project of Jiangxi Provincial Education Department (60224), Key Projects of Jiangxi Natural Science Foundation (20171ACB20003), and Key Research and Development Projects of Jiangxi Natural Science Foundation (20192BBG70067).

## Author contributions

Lingbing Zeng and Xiaotian Huang conceived and designed the experiments; Hongxia Zhang, Lingbing Zeng, Qiong Liu, Guilin Jin, Jieyu Zhang and Zengbin Li performed the experiments. Hongxia Zhang and Qiong Liu analyzed the data; Hongxia Zhang, Lingbing Zeng and Xiaotian Huang wrote the manuscript.

## Conflicts of interest

The authors declare that they do not have any conflicts of interest.

## Figure legends

**Supplemental Figure 1.** CVB3 VP1 results in a specific cytopathic effect to HPAC cells. Cell adherence was observed at 48 h after transfection or infection through the comparative analysis of bright-field images. Scale bar = 100 µm.

**Supplemental Figure 2.** The expression levels of VP1 increased in a time-dependent. Western blot analysis of VP1 expression in Mock, pBud, pBud-VP1 and CVB3 groups.

**Supplemental Figure 3.** CVB3 VP1 inhibits cell proliferation in HeLa cells. **(A)** HeLa cells were transfected with pBud-VP1 and pBud plasmid, and the cell proliferations were separately evaluated using a CCK-8 assay. **(B)** The activity of DNA replication was examined using EdU incorporation in HeLa cells. **(C)** The number of adherent HeLa cells in different groups was counted using Image Pro Plus (IPP) software. Mean ± range of values for the counts of cell adherence in replicate experiments.

**Supplemental Figure 4.** CVB3 infection induces G1/S phase accumulation of HPAC and HeLa cells. After HPAC and HeLa cells were treated with thymidine again, HPAC **(A)** and HeLa **(B)** cells were mock infected or infected with CVB3 at a MOI of 5. These cells were released from the thymidine block and collected according to the indicated release time (0, 6 and 9 or 12 hrs); the cells were analyzed by flow cytometry. The percentage of cells in each phase of the cell cycle is showed as mean ± SEM of three independent experiments. (\**P* < 0.05, \*\**P* < 0.01).

**Supplemental Figure 5.** Other structural proteins of CVB3 cannot arrest the cell cycle at the G1/S phase. The structural proteins of CVB3 VP2, VP3 and VP4 infected double-thymidine synchronized cells, with GFP as a control. These cells were then released and collected according to the indicated release time (0, 3, 6, or 9 hrs). The percentage of cells in each phase of the cell cycle is shown as mean ± SEM of three independent experiments. (\**P* < 0.05, \*\**P* < 0.01).

**Supplemental Figure 6.** MAT1 and VP1 intracellular localization in CVB3 infected HeLa cells. Representative confocal immunofluorescence microscopic images of MAT1 and VP1 stained with rabbit anti-MAT1 (green) and mouse anti-VP1 antibodies (red); the nuclei are labeled with DAPI. Scale bar = 10 µm.

**Supplemental Figure 7.** Confocal microscopy analysis of the abundance of VP1 and MAT1 in HeLa cells. Cells were detected with monoclonal antibodies to MAT1 (Alexa-488) and polyclonal anti-VP1 (fluorescein; red), and counterstained with DAPI to show the nucleus. The MAT1 and VP1 images were merged. Scale bar: 10 µm.

**Supplemental Figure 8.** Immunoblot analysis of the cytoplasmic-localized accumulation of MAT1, CDK7, Cyclin H in CVB3 infected, pBud-VP1 and pBud transfected cells by specific antibodies.

